# Simultaneous multi-site editing of individual genomes using retron arrays

**DOI:** 10.1101/2023.07.17.549397

**Authors:** Alejandro González-Delgado, Santiago C. Lopez, Matías Rojas-Montero, Chloe B. Fishman, Seth L. Shipman

## Abstract

Our understanding of genomics is limited by the scale of our genomic technologies. While libraries of genomic manipulations scaffolded on CRISPR gRNAs have been transformative, these existing approaches are typically multiplexed across genomes. Yet much of the complexity of real genomes is encoded within a genome across sites. Unfortunately, building cells with multiple, non-adjacent precise mutations remains a laborious cycle of editing, isolating an edited cell, and editing again. Here, we describe a technology for precisely modifying multiple sites on a single genome simultaneously. This technology – termed a multitron – is built from a heavily modified retron, in which multiple donor-encoding msds are produced from a single transcript. The multitron architecture is compatible with both recombineering in prokaryotic cells and CRISPR editing in eukaryotic cells. We demonstrate applications for this approach in molecular recording, genetic element minimization, and metabolic engineering.

## INTRODUCTION

Multiplexing – the act of consolidating multiple discrete elements into a single composite channel – has enabled genomic technologies to scale toward the complexity of the biology we hope to understand. Today, one might use multiplexed DNA synthesis to make a library of distinct CRISPR gRNAs on a single synthesis chip, then use multiplexed experimental design to clone and transfect that library of gRNAs across cells in a single culture, and finally use multiplexed sequencing to analyze the effect of the perturbation on a single sequencing flow-cell^1, 2^. This now-standard multiplexed gRNA workflow has allowed scientists run experiments across every gene in parallel with barely more effort than they might have previously put into determining the effect of single gene. However, the typical multiplexing of a gRNA library precludes an important level of analysis: it is implemented across cells, where a single edit is made per genome, and thus cannot be used to study the interaction of mutations within a genome.

Technologies for multiplexing *within* genomes – where multiple distinct, non-adjacent edits are made using a single, consolidated editor – are much more limited. Yet, applications for multiplexing within genomes abound in both fundamental biology (e.g. studying epistasis, long-range gene regulation, and genome organization) and biotechnology (e.g. metabolic engineering, molecular recording, and genome minimization). These complex applications require precise mutations, not genomic scars or transcriptional perturbations. Precision is essential to understand combinatorial genome complexity, such as probing compensatory mutations across genes in a complex or interrogating enhancer-promoter interactions, and is necessary to build nuanced technological advances, such as ribosome-dependent tuning of gene expression in a metabolic pathway.

In bacteria, the most commonly used approach to introduce combinatorial, precise mutations is MAGE (multiplexed automated genome engineering), which relies on single stranded DNA (ssDNA) recombineering^3–5^. A eukaryotic version of this technology has been developed to extend this approach to yeast^6^. However, MAGE is limited by its requirement for numerous labor-intensive recombineering cycles required to attain efficient combinatorial editing rates, and by its reliance on exogenously-delivered oligonucleotides that leave no trackable plasmid element for phenotyping by proxy^7^. Base-editing (BE) and prime-editing (PE)^8–10^ are two other precise editing approaches that can be multiplexed^11–16^. Base-editors are the simplest to multiplex using tandem gRNAs, but are limited to single base mutations of a defined type (either A•T-to-G•C or C•G-to-T•A)^11, 13–14^. Prime-editors have also been multiplexed, but the complexity of the editing elements grows quickly with additional sites. In bacteria, multiplexed prime editing requires a three plasmid system, and multiple edits occur on the same genome in less than 1% of cells^12^, while systems built for human and plant cells require two gRNAs per site in addition to the editing template, which can create issues with the assembly of multiplexed plasmids^14–16^.

Another way to introduce precise mutations that is compatible with both prokaryotic and eukaryotic editing is to produce editing donors inside a cell using modified retrons. Retrons are bacterial tripartite systems that have been shown to provide phage defense^17–20^. Two of the components of the retron operon are a reverse transcriptase and a small (200-300 base), structured non-coding RNA (ncRNA). The reverse transcriptase recognizes and partially reverse transcribes the ncRNA into a single-stranded DNA fragment that is present at the abundance of a cellular transcript^21–24^.

We and others have previously shown that the retron ncRNA can be modified to encode an editing donor to precisely edit the genomes of bacteria, phage, plant, yeast, and even human cells^25–31^. However, these retron-derived editors have only been used to edit genomic positions one at a time. Here, we describe a substantial modification of the retron ncRNA to produce multiple editing donors simultaneously from a single transcript after reverse transcription. We show that these multiplexed, arrayed retron elements – termed multitrons – can be paired with single-stranded annealing proteins to edit prokaryotic genomes and with CRISPR components to edit eukaryotic genomes^26–29^. We demonstrate utility with proof-of-concept applications in molecular recording, multiplexed deletions, and metabolic engineering.

## RESULTS

### Multiplexed recombineering from multiple donors in a retron msd

The use of retrons in bacterial recombineering was originally developed for applications in molecular recording^25^, and has more recently been optimized to install single targeted edits and interrogate biology^26–29^. To do so, a retron ncRNA – which can be divided into two regions: an msr (multicopy single-stranded RNA) that is not reverse transcribed and an msd (multicopy single-stranded DNA) that is reverse transcribed – is modified to encode an editing donor within the msd region. This modified ncRNA is expressed in cells along with a retron reverse transcriptase (e.g. retron Eco1-RT) that reverse transcribes the retron msd to produce an editing donor (RT-Donor). An overexpressed single-stranded annealing protein (SSAP, e.g. CspRecT) and the host single-stranded binding protein (SSB) promote annealing of the RT-Donor to the lagging strand of a replicating chromosome to install the edited sequence^32–33^.

We aimed to further modify retrons to create multitron editors, capable of multiplexed editing of a single genome from a consolidated retron element generating multiple RT-Donors per transcript. Recombineering via oligonucleotide donors is most efficient with donors between 70 and 90 bases long^3^, which is also the ideal range for retron recombineering donors^28, 31^. Yet, retron RTs are capable of reverse transcribing much longer RT-Donors, even up to an entire gene length^26^. Thus, we initially tested a multitron architecture that encodes multiple 70 bp donors end-to-end within a single msd loop (**Fig 1a**) using the two tandem donors to make point mutations in both the *rpoB* and *gyrA* genes in *E. coli*. We tested two versions of this multitron with the donors in each of the possible orders in the msd as well as a control *rpoB* singleplex editor. Both tandem multitron variants edited both sites, and editing rates for *rpoB* were comparable in the singleplex versus multitron configurations (**Fig 1b**).

**Figure 1.**
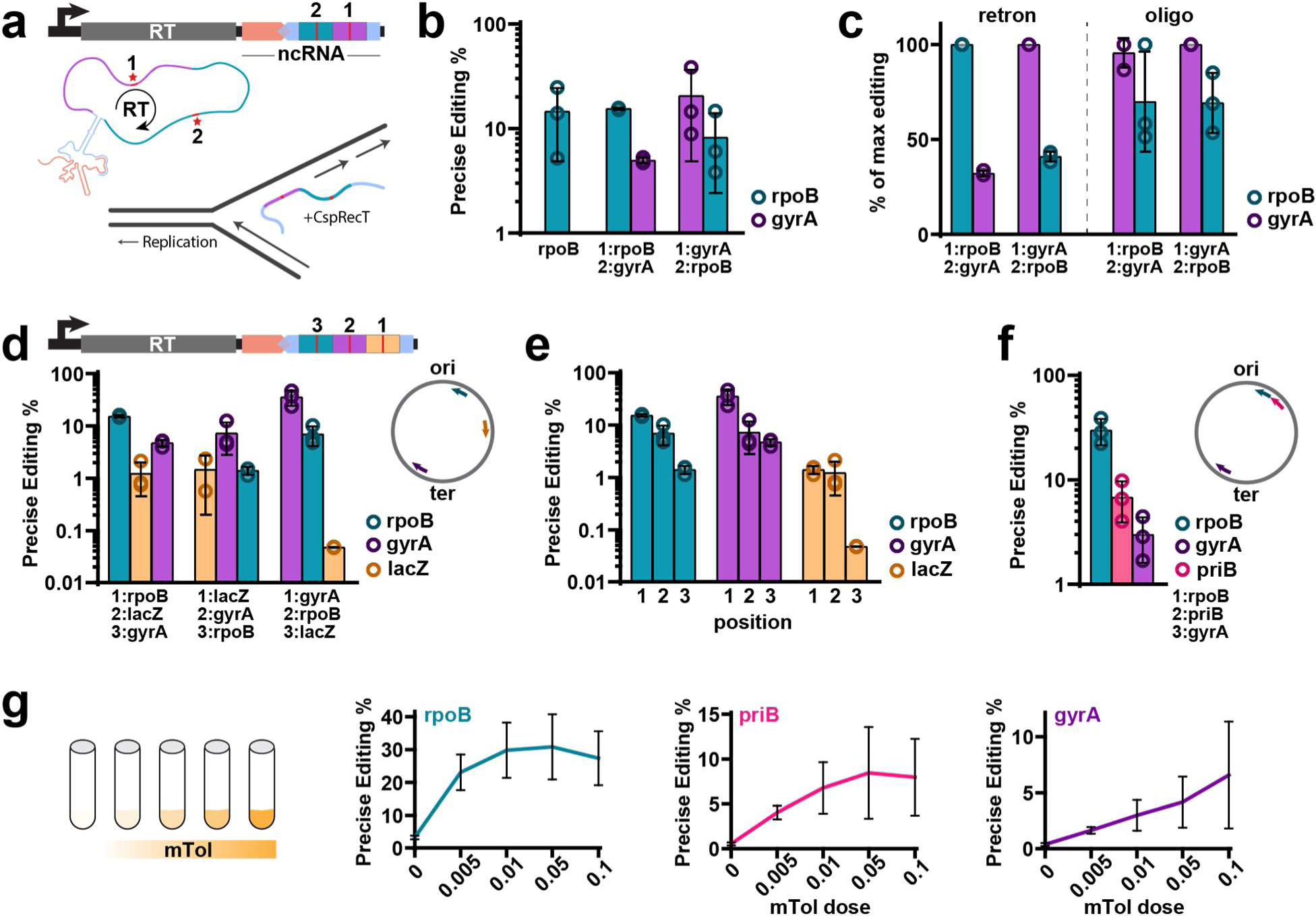
Encoding several donors in a retron msd enables multiplexed retron recombineering in bacteria. **a.** Top: schematic of the retron recombineering operon with two donors encoded within the msd region (light blue). Donor labels 1 (purple) and 2 (green) indicate the order in which the donor is reverse transcribed by the retron RT (grey). Bottom: schematic of the retron recombineering process, which occurs during replication in the lagging strand with the participation of CspRecT protein. Red stars represent the desired mutations to be integrated in the bacterial chromosome. **b.** Quantification of precise editing rates of the *rpoB* locus alone and both *rpoB* and *gyrA* loci in bacteria. The order in which the donors are reverse transcribed (1 or 2) is indicated for each condition. For b, c, d, e, and f, data were quantified by Illumina sequencing after 24h of editing, circles show each of the three biological replicates, bars are mean ±SD (one-way ANOVA, effect of condition on *rpoB* editing *P*=0.2616). **c**. Comparison of donor order for retron-encoded donors versus donors encoded on an oligonucleotide matching the retron RT-DNA. Editing is shown as percent of maximum precise editing for each condition, illustrating that the retron is influenced by position effects that are not found when using an oligonucleotide donor (one-way ANOVA effect of conditions *P*<0.001; Tukey’s corrected effect of retron order *P*<0.0001, oligo order *P*=0.9842). **d**. Top: schematic of the retron recombineering cassette with 3 donors encoded in the msd region. The numbers above the donors indicate the order of reverse transcription. Bottom: quantification of precise editing rates of bacterial *rpoB*, *gyrA*, and *lacZ* loci. Right: schematic of the bacterial chromosome indicating donor position and strand with respect to the origin of replication (lagging strand for *rpoB* and *gyrA* donors and leading strand for the *lacZ* donor). **e.** Replot of the data in d, illustrating the effect of position on editing at each site (two-way ANOVA effect of position *P*<0.0001). **f.** Quantification of precise editing rates for *rpoB*, *gyrA* and *priB* loci in bacteria, in the same architecture shown in d. Right: schematic of bacterial chromosome indicating donor position and strand respect to the origin of replication. In this case, all donors are in the lagging strand. (one-way ANOVA, effect of editing site *P*=0.0015). **g**. Use of multiplexed retron recombineering to improve analog molecular recording technologies. (left) Increasing amounts of m-toluic acid (mTol) are recorded using a retron-derived analog recorder; (right) quantification of precise editing rates for *rpoB*, *gyrA* and *priB* loci using different amounts of mTol. Error bars indicate the standard deviation for three independent biological replicates. Additional statistical details in **Supplemental Table 1**.

When comparing the two multitron versions, we noticed that the site edited by the first donor in the multitron tended to have a higher editing rate than the site edited by the second donor. The donor in position one is reverse transcribed first, so the editing difference could be due to a small effect of RT processivity, or due to a positional effect of the donors after reverse transcription. To distinguish between these possibilities, we compared the relative editing efficiencies at each site using the multitrons versus synthetic oligonucleotides of the same sequence as the tandem RT-Donors. Unlike RT-Donors produced by multitrons, oligonucleotide donors had similar relative editing rates across the sites independent of their donor position (**Fig 1c**), consistent with an effect of RT processivity.

We next tested three donor multitrons in the tandem msd architecture, using a third donor targeting *lacZ* on the leading strand (less effective than targeting the lagging strand). All three sites were edited in each of the three permutations of donor order (**Fig 1d**), with the same positional bias for higher editing at the 5’ end of the RT-Donor (**Fig 1e**). Although the positional bias is a bug in our intended design, we wondered whether it could be exploited to create a range of editing efficiencies for analog molecular recording. Retrons have previously been used as analog molecular recorders capable of detecting the magnitude and duration of a specific input by accumulating precise mutations in the genome^25^. These analog molecular recorders are, however, limited to operating in the linear range of the interaction between reporter and editing efficacy. We reasoned that using a tandem multitron could add robustness by expanding the dynamic range of a recording across multiple sites. We constructed another multitron encoding three lagging donors (*gyrA*, *priB*, *rpoB*) driven by an m-toluic acid (mTol)-inducible promoter. Here too, we found that the editing rates were inversely proportional to the order of donor reverse transcription at maximal induction (**Fig 1f**). As a result, the editing rates for each site saturate at different mTol concentrations when used as an analog recorder of mTol (**Fig 1g**), effectively increasing the dynamic range of the recorder.

### Improved Multiplexed Editing Using Donors in Retron Arrays

To overcome the effect of donor position inside a single msd loop, we engineered a different version of the multitron architecture composed of an ncRNA array with multiple msr-msd regions in tandem, each one containing a distinct donor to edit a unique target site (**Fig 2a**). With this arrayed ncRNA multitron, the retron RT has different substrates available within a transcript to generate multiple RT-donors independently, each at the same distance from an internal RT priming site. We tested the ability of this arrayed ncRNA multitron to edit *rpoB* and *gyrA* versus singleplex retron editors, and found that the arrayed ncRNA multitron performed as well or better than the singleplex versions (**Fig 2a**). However, this arrayed ncRNA created a new constraint. The length of the ncRNA donor unit is 229 bp and the arrayed design adds 109 bp of direct repeat for each additional editor due to msr duplication, both of which pose challenges for the synthesis and assembly of new multitron plasmids.

**Figure 2.**
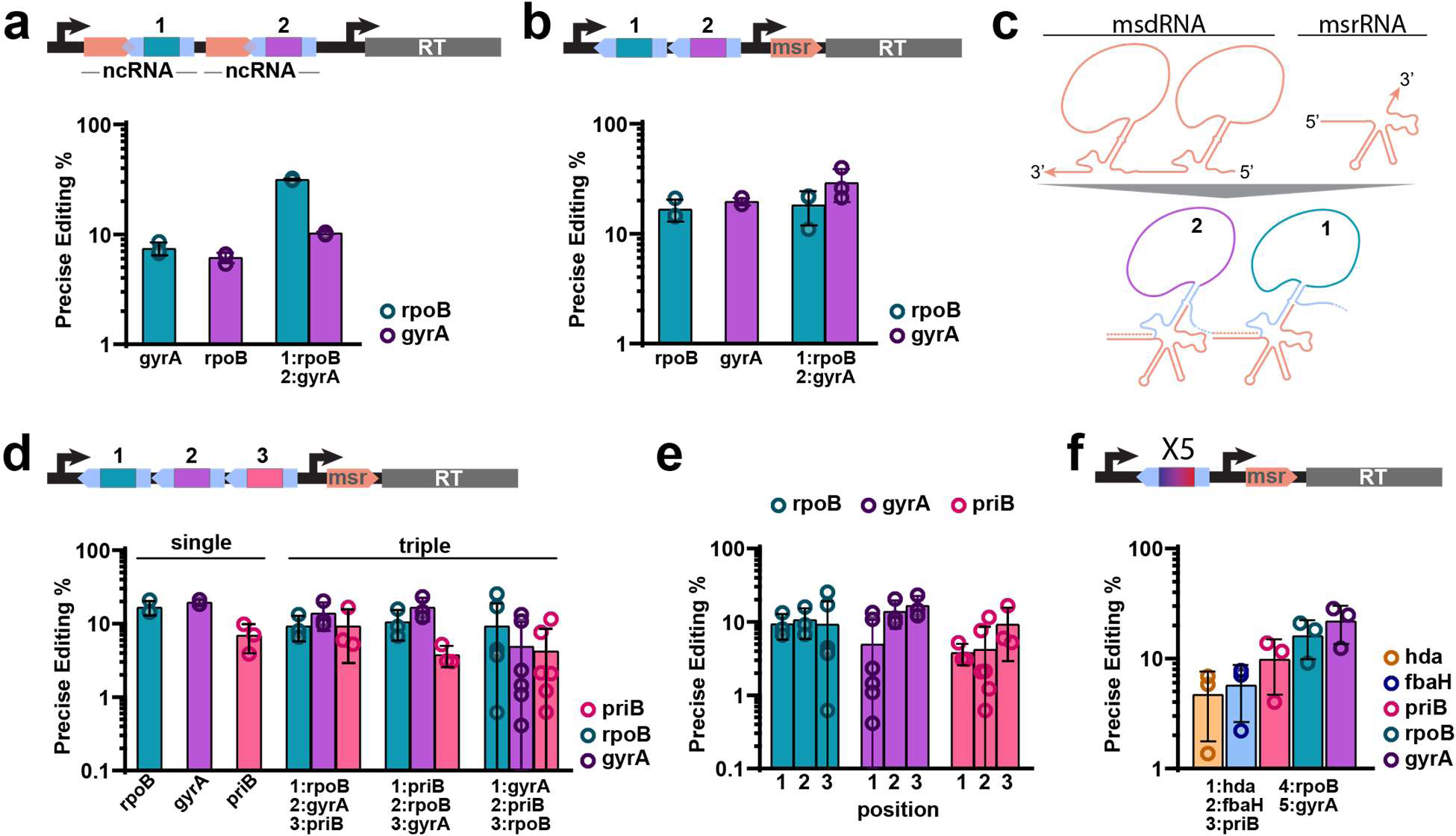
Improved multiplexed editing using donors in arrayed retron msds. **a.** Top: schematic of retron recombineering using 2 independent ncRNAs. Each msd region (blue) encodes a different donor (1 and 2). Bottom: quantification of precise editing rates for precise editing of *gyrA* or *rpoB* alone or simultaneously (unpaired t-test, singleplex versus multiplex, *rpoB P*<0.0001, *gyrA P*=0.0006). **b.** Top: Schematic of retron recombineering using an msd array with a single msr sequence in trans. Bottom: quantification of precise editing rates for precise editing of *rpoB* or *gyrA* alone or simultaneously (unpaired t-test, singleplex versus multiplex, *rpoB P*=0.7312, *gyrA P*=0.1702). **c**. Top: schematic of arrayed msd and msr transcription products. Arrayed msd is transcribed as a single transcript. Bottom: schematic of RT-DNA production using as template an arrayed msd. 1 and 2 indicates the number of the msd in the arrayed msd. **d**. Top: schematic of 3x arrayed msd. Bottom: quantification of precise editing of *rpoB*, *gyrA* or *priB* edits alone or simultaneously. **e.** Replot of the data in d, illustrating the effect of position on editing at each site (two-way ANOVA, effect of position *P*=0.1138). **f.** Quantification of precise editing using a 5x arrayed msd to edit *hda*, *fbaH*, *priB*, *rpoB* and *gyrA*. Data in a, b, d, e and f were quantified by Illumina sequencing after 24h of editing, circles show each of the three biological replicates, bars are mean ±SD. Additional statistical details in **Supplemental Table 1**.

Therefore, we engineered a third multitron version composed of an msd array rather than an ncRNA array. In this case, each msd encodes a distinct donor as in the previous version, but the msr is expressed in trans as a separate transcript (**Fig 2b**). This trans msr arrangement was previously shown to be a tolerated modification for reverse transcription of endogenous retron msds^34^. In practice, this reduces the editing unit to 149 bp and reduces the length of the longest direct repeat to 74 bases. The trans msr can interact with any of the arrayed msds, again keeping the donor at a constant distance from the site of RT priming (**Fig 2c**).

We tested editing by the arrayed msd multitron versus singleplex editors and found no difference in editing rates at either site (**Fig 2b**). The trans msr arrangement in fact yielded consistently higher editing rates than the endogenous retron ncRNA architecture in both singleplex and multiplexed forms throughout this project. Although the msd array and msr/RT transcript contain no terminator between them and could potentially be transcribed as a single unit rather than the intended trans arrangement, we found both sites could be edited at a similar efficiency when using a plasmid containing a terminator between the msd array and the msr (**Supplementary Fig 1**).

To test whether donor position inside the msd array multitron affects editing, we constructed three multitron variants with donors to edit *priB*, *rpoB* and *gyrA* genes in each possible order. All three sites were edited by each multitron variant (**Fig 2d**), and there was no effect of donor position using arrayed msds (**Fig 2e**). Finally, to push the limits of within-genome multiplexing, we constructed an arrayed msd multitron to simultaneously edit 5 target sites (*hda*, *fbaH*, *priB*, *rpoB* and *gyrA*). Editing rates ranged from 5 to 25% for each site, illustrating that arrayed msd multitrons are a potent tool for multiplexed genome editing technologies (**Fig 2f**).

### Increasing Limits of Deletion Size Using Nested Multitrons

One benefit of using retron-derived donors is that they support a broad range of precise mutations, including insertions, deletions and replacements. However, when recombineering with either retron RT-Donor or oligonucleotide donors, the efficiency of inserting and deleting base pairs is inversely related to the size of the edit^3, 31^. This is presumably intrinsic to the mechanism of recombineering, a result that we replicated here using RT-Donors to delete 1 to 100 bp, finding a declining efficiency with deletion size whether using an endogenous ncRNA architecture or the trans msr architecture (**Fig 3a**).

**Figure 3.**
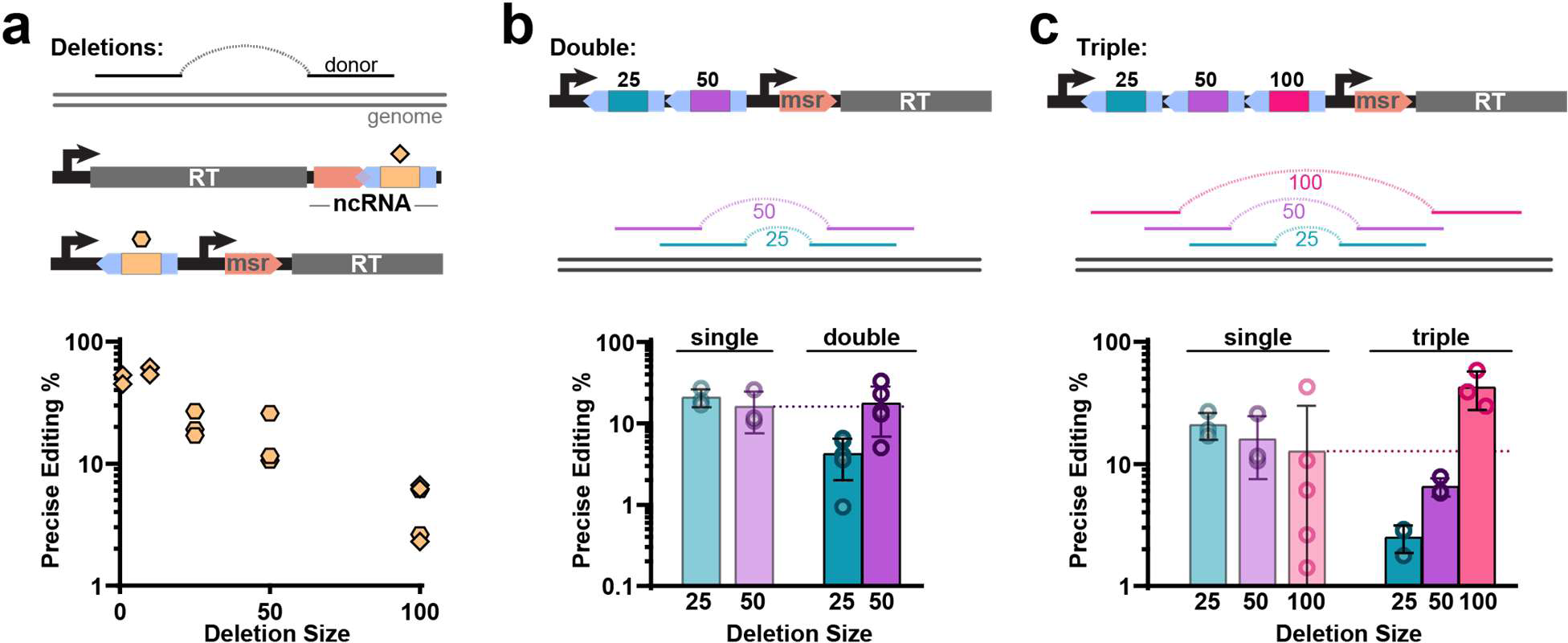
Increasing limits of deletion size using nested deletion donor arrays. **A.** Top: Schematic of genome deletions using retron recombineering. Middle: schematic of a standard retron cassette to make deletions (top) with the donor represented by a diamond and an arrayed msd retron cassette with the donor represented by a hexagon. Bottom: quantification of precise editing rates for a single deletion of 1 bp, 10 bp, 25 bp, 50 bp or 100 bp deletions by Illumina sequencing after 24h of editing. Diamonds show deletions from a standard architecture and hexagons show deletions using an arrayed architecture (one-way ANOVA, effect of deletion size *P*<0.0001). **b.** Top: Schematic of arrayed msd retron cassette with two donors to make 25 and 50 bp deletions. Middle: Schematic of a nested deletion strategy using two donors to delete 25 bp and 50 bp. If the 25 bp occurs first, the 50 bp deletion becomes a 25 bp deletion. Bottom: Quantification of precise editing rates for single 25 and 50 bp deletions, and for the nested 50 bp deletion. **C.** Top: Schematic of arrayed msd retron cassette with three donors to make 25, 50 bp and 100 bp deletions. Middle: Schematic of a nested deletion strategy using three donors to delete 25 bp, 50 bp and 100 bp. Bottom: Quantification of precise editing rates for single 25 bp, 50 bp and 100 bp deletions, and for each deletion using the nested strategy (unpaired t-test, singleplex versus multiplex 100bp deletion, *P*=0.0485). Data in b and c were quantified by Illumina sequencing 24h after of editing, circles show each of the three biological replicates, bars are mean ±SD. Additional statistical details in **Supplemental Table 1**.

We wondered whether we could overcome this limitation on deletion efficiency at larger sizes by using arrayed msd multitrons encoding a series of nested deletion donors. A nested deletion series consists of multiple donors intended to make deletions of increasing size progressively at same locus. If the smallest deletion succeeds, it creates a smaller target size for a previously disfavored large deletion. We explored nested deletions by first comparing the editing efficiency of single 25 and 50 bp deletions in the *lacZ* gene with simultaneous deletions of overlapping 25 and 50 bp at the same location using a multitron (**Fig 3b**). The 50 bp deletion was not significantly less efficient than the 25 bp deletion using singleplex retron donors so, unsurprisingly, the rate of 50 bp deletions by the multitron version was not significantly increased. However, the rate of the 25 bp deletion was decreased by the multitron, suggesting that 25 bp deletions were being converted into 50 bp deletions.

Next, we tested a multitron containing a 25, 50, and 100 bp nested deletion donor series (**Fig 3c**). In this case, the previously disfavored 100 bp deletion was significantly more efficient using the multitron series than using the singleplex deletion donor. In fact, this strategy created a 100 bp deletion in ∼42% of genomes, overcoming an intrinsic inefficiency in recombineering deletions. Furthermore, the multitrons generated a heterogeneous population of genetic elements with different deletions sizes that could be used to probe functional domains of a target gene or miniature versions of a protein of interest.

### Metabolic Engineering in Bacterial Genomes Using Multitrons

We next pushed toward a proof-of-concept use of multitrons in metabolic engineering, which clarified the last steps of system optimization. First, the five molecular elements required for multitron recombineering – msd array, msr, RT, RecT, and dominant negative mutL (to suppress mismatch repair for single base mutations) – which had been spread across two plasmids were cloned into a single plasmid to simplify the overall approach (**Supplementary Fig 2a**). However, editing rates for the *rpoB* gene were ∼5x lower using the single plasmid compared to the previous two plasmid system (∼5% and ∼25%, respectively). To increase recombineering efficiency, we added an *E. coli* optimized ribosome binding site (RBS) immediately upstream of only the RT gene or both the RT and the CspRecT genes, both of which increased editing rates but still fell short of the level achieved by the two-plasmid system (**Supplementary Fig 2a**).

We next changed the origin of replication for the single plasmid system, opting for a temperature-sensitive origin so that the plasmid becomes curable after editing by moving from a permissive temperature (30°C) to a non-permissive temperature (37°C)^35, 36^. Interestingly, the editing rates using this single plasmid finally reached comparable levels to those of the previous the two-plasmid system (**Supplementary Fig 2a**). Finally, we titrated the inducer concentration to reach optimal editing at 1% arabinose (**Supplementary Fig 2b**).

We next assessed the ability of re-optimized, temperature-sensitive arrayed msd multitrons to simultaneously edit five positions (*hda, fbaH, priB, rpoB* and *gyrA*). All sites were precisely edited after 24h, and editing continued to increase over the next 24h following a passage, illustrating the continuous nature of the retron-derived editing (**Fig 4a**).

**Figure 4.**
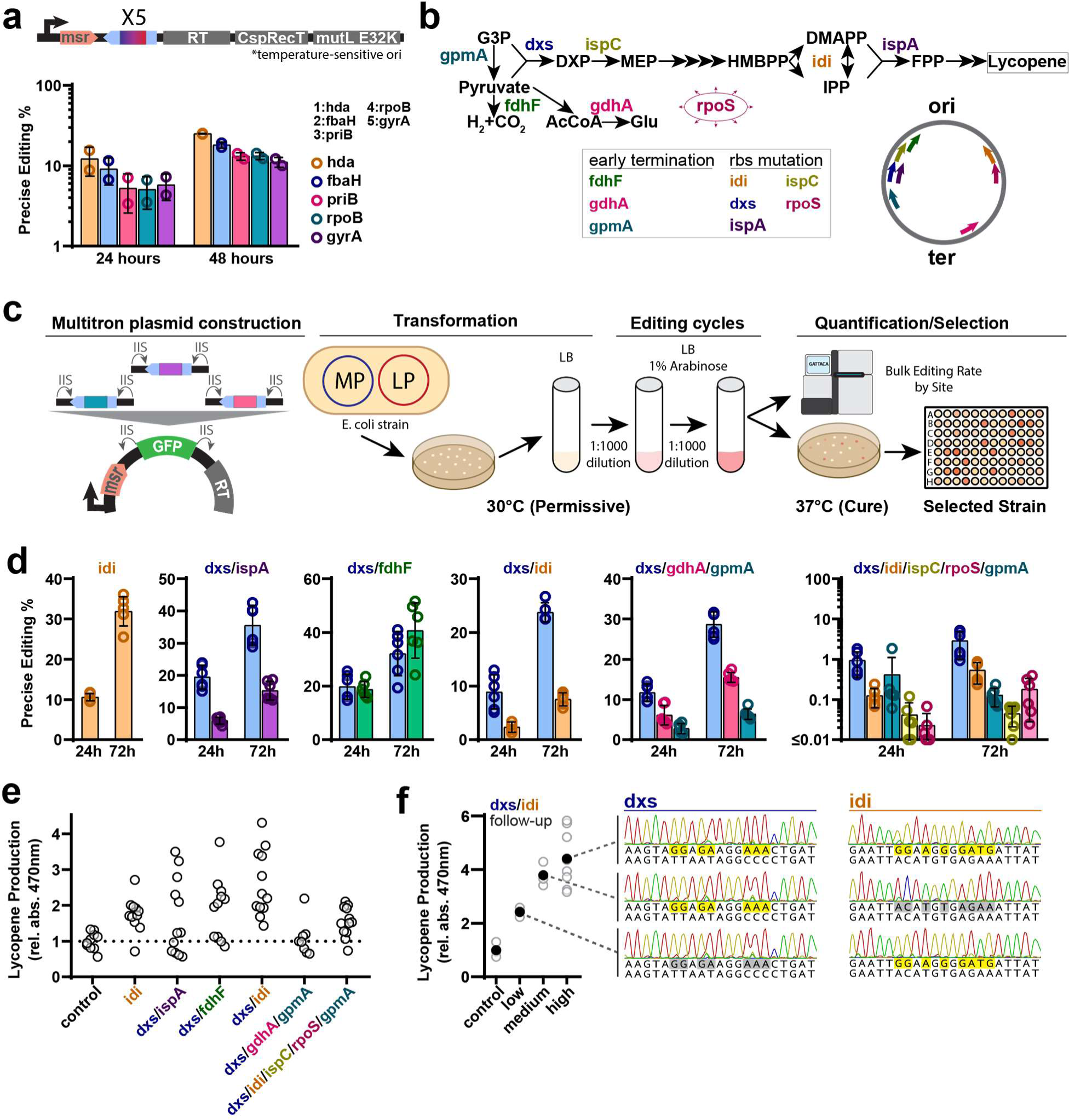
Metabolic engineering using multitrons. **a.** Top: architecture of the multiplexed retron recombineering cassette in the temperature sensitive plasmid. The operon is composed of a single msr followed by 5x arrayed msds with donors and the genes encoding the RT, the CspRecT and the dominant negative MutLE32K. Bottom: quantification of precise editing rates using a 5x arrayed msd to edit *hda*, *fbaH*, *priB*, *rpoB* and *gyrA* by Illumina sequencing 24h and 48h after of editing (two-way ANOVA, effect of expression time *P*<0.0001). Circles show each of the three biological replicates, bars are mean ±SD. The order of the donors in the arrayed msd is indicated. **b.** Top: Schematic of the lycopene biosynthesis pathway, with key genes to increase lycopene production highlighted. Bottom: Schematic of metabolic engineering of lycopene biosynthesis pathway using multiplexed retron recombineering. **c.** The donors are cloned into a temperature sensitive backbone using a golden gate assembly protocol. Single colonies are grown for 24 h and then induced with arabinose. This cycle is repeated by making 1:1000 dilutions for several days. Editing rates are measured by Illumina sequencing and cultures are plated to select individual colonies based on color for quantification of lycopene production. **d.** Quantification of precise editing rates using different recombitron plasmids containing a variable number of donors to edit genes in the lycopene pathway, quantified by Illumina sequencing after 24h and 72h. Circles show each of the six biological replicates, bars are mean ±SD. **e.** Quantification of lycopene production in single colonies. Lycopene production was normalized against the average production of the control, which contains the pAC-LYC but was not exposed to the recombineering process. Each point represents a colony (n=12). **f.** Quantification of lycopene production from colonies re-isolated from samples in the low (∼2X control), medium (∼3X control), and high (∼4x control) production clusters of the *dxs/idi* condition. Open circles are individual colony values and closed circles are the mean. Sanger sequencing examples to the right illustrate the genotype of each subset (all individual colonies within a condition have identical genotypes). Additional statistical details in **Supplemental Table 1**.

To test multitrons in the context of metabolic engineering, we chose to focus on increasing production of lycopene by modifying genes in its biosynthetic pathway (**Fig 4b**). We selected eight bacterial genes which have been shown to affect lycopene yield^3, 37–39^ (**Fig 4b**). Five of them (*dxs, idi, ispA, ispC, rpoS*) were subjected to modification of their RBS regions to enhance their similarity to the canonical Shine-Dalgarno sequence (TAAGGAGGT)^40^. The other three genes (*gmpA, gdhA, fdhF*) were specifically targeted for inactivation by the introduction of premature stop codons within their open reading frames.

We established a general workflow for metabolic engineering using multitrons (**Fig 4c**; **Material and Methods**). The multitron plasmid (MP) was generated using a one-pot golden gate approach^41^ to clone arrayed msds encoding different donors. The MP was next transformed into the bacterial host harboring the lycopene plasmid (LP, a plasmid containing three essential genes (*crtE, crtI, crTB*) required for lycopene production^42^. Editing cycles were carried out at the permissive temperature (30°C), with dilutions of the culture after every cycle. Editing targets were sequenced in bulk using Illumina MiSeq to determine overall efficiencies. In parallel, cells were plated at 37°C to cure the MP. Finally, red colonies (indicative of lycopene) from the plates were selected for further quantification of lycopene production levels (**Fig 4c**).

In total, we tested six different arrayed msd multitrons across this workflow, containing target gene donors in combinations that have been have been shown to increase lycopene yield^3^. Editing rates were measured after cycles 1 and 3 of editing (24h and 72h, respectively) showing values that increase with time (**Fig. 4d**). After 72h of editing, the precise editing rates when making one or two mutations ranged from 10 to 40%. When making three or five mutations, editing rates were lower, which could be due to the known negative fitness effect^3^ of these mutations on the bacterial growth (**Fig 4d**).

We measured relative lycopene production from 84 isolated red colonies after plating cultures on LB agar plates after editing (**Fig 4e**). In each case other than the control, individual colonies produced variable amounts of lycopene, likely resulting from the intended genotypic diversity generated by the editing. As an example, the most productive isolate after RBS optimization of *dxs* and *idi* genes increased lycopene production by more than 400% of control values, there was a second production cluster around 300% of control, and a final cluster around 200% of control (**Fig 4e**). We reasoned that these three different clusters may represent a single *dxs* mutation, a single *idi* mutation, and both together. To test that hypothesis, a representative of each cluster was selected and re-streaked for colonies, which were re-measured for lycopene and Sanger sequenced. Indeed, that the best producing isolate carried RBS mutations of both *dxs* and *idi* genes, second-best had only the *dxs* mutation, and the third-best had only the *idi* mutation (**Fig 4f**). Thus, a multiplexed experiment generates a diversity of genotypes and corresponding phenotypes across multiple sites simultaneously.

### Multitrons with CRISPR Editing in Eukaryotic Cells

Given the success of the arrayed msd multitron in recombineering, we next sought to expand the utility of this technology to eukaryotic cells. Retron RT-Donors have been used in *S. cerevisiae* in combination with CRISPR Cas9 and gRNAs to install precise mutations via templated repair of a cut site (**Fig 5a**). The architecture of the donor element in yeast is typically a retron ncRNA fused to a CRISPR gRNA and scaffold, all surrounded by ribozymes to excise the editing elements from an mRNA. Given the goal of engineering a eukaryotic msd array, the relatively large, structured ribozymes present a potential engineering hurdle if they need to be multiply duplicated. Therefore, we first tested replacement of the ribozymes with Csy4 recognition sites and Csy4 nuclease expression by comparing a singleplex retron-derived precise editor of the *ADE2* locus in the standard arrangement using ribozymes against an alternate version in which the flanking ribozymes were replaced by Csy4 recognition sites. In both cases, we tested editing with or without the inclusion of a Csy4 gene in an integrated, inducible, genomic cassette that also expresses the retron RT and Cas9. We found, as expected, no effect of Csy4 expression on the ribozyme version of the precise editor, but a dramatic effect of Csy4 expression on the alternate version with Csy4 sites. Precise editing nearly matched the efficiently of the ribozyme version with Csy4 expression, but was sharply reduced in its absence, indicating that processing of the non-coding elements is required and can be achieved using Csy4 (**Fig 5a**).

**Figure 5.**
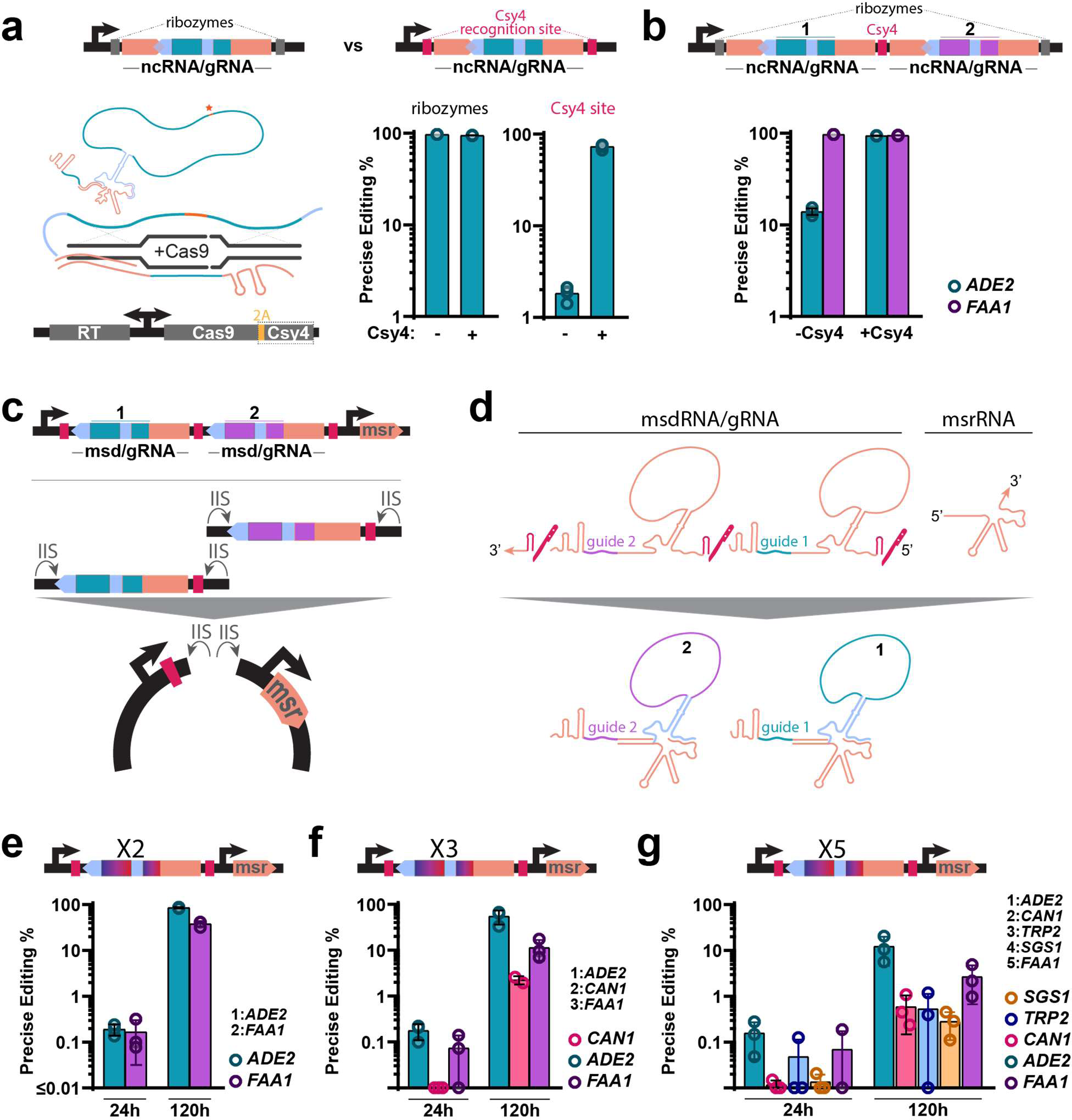
Arrayed retron msds enable multiplexed editing in yeast cells. **a.** Top: Schematic of the donor encoding retron ncRNA/gRNA expression cassette expressed from a Gal7 Pol II promoter and flanked by ribozymes versus a new construction replacing ribozymes with Csy4 sequences. Bottom left: schematic of a retron ncRNA-Cas9 gRNA hybrid for genome editing in yeast, depicted above the protein-coding expression cassette which is inserted into the yeast genome. Bottom right: quantification of precise editing of the *ADE2* locus in yeast by Illumina sequencing after 48h of editing. Circles show each of the three biological replicates, bars are mean ±SD; absence/presence of Csy4 in the protein-coding expression cassette is shown below the graph (Sidak’s corrected multiple comparisons, effect of Csy4 expression, ribozyme construction *P*=0.2779, csy4 construction *P*<0.0001). **b.** Top: schematic of an arrayed retron ncRNA-Cas9 gRNA expression cassette, expressed from a Gal7 Pol II promoter, flanked by ribozymes, and separated by a Csy4 sequence. The retron editors in positions 1 and 2 target the *ADE2* and *FAA1* locus, respectively. Bottom: quantification of precise editing of the *ADE2* and *FAA1* loci in yeast by Illumina sequencing after 48h of editing. Circles show each of the three biological replicates, bars are mean ±SD; absence/presence of Csy4 in the protein-coding expression cassette is shown below the graph (Sidak’s corrected multiple comparisons, effect of csy4 expression, ADE2 *P*<0.0001, FAA1 *P*=0.0012). **c.** Top: schematic of an arrayed retron msdRNA-Cas9 gRNA expression cassette, expressed from a Gal7 Pol II promoter, flanked and separated by a Csy4 sequence; the msrRNA is expressed in trans from a SNR52 Pol III promoter. Bottom: assembly schematic for one-pot Golden Gate cloning of multiple msdRNA-sgRNA editors. **d**. Schematic showing the presumed processing, annealing and reverse-transcription involved in the generation of editing donors from arrayed retron msdRNA-Cas9 gRNA cassettes. **e-g**, top: schematic of 2x, 3x or 5x arrayed retron msdRNA-Cas9 gRNA expression cassettes, as shown in c. Bottom: quantification of precise editing of the various yeast loci targeted by the retron editors shown above, by Illumina sequencing, after 24 and 120h of editing. The editors target *ADE2* and *FAA1* (e); *ADE2*, *CAN1* and *FAA1* (f); and *ADE2*, *CAN1*, *TRP2*, *SGS1* and *FAA1* (g). Two-way ANOVA, effect of expression time, e *P*<0.0001, f *P*<0.0001, g *P*=0.0038. Circles show each 3 biological replicates, bars are mean ±SD. Additional statistical details in **Supplemental Table 1**.

We next tested a eukaryotic multitron based on an array of ncRNA/gRNAs targeting *ADE2* and *FAA1* for precise mutations of three base pairs each. For each site, the ncRNA encoding the donor for the site was fused to the gRNA for the same site. The two sites were separated by a Csy4 recognition site and the double ncRNA/gRNA array was surrounded by ribozymes (**Fig 5b**). Both sites were edited to nearly 100% in the presence of Csy4 expression. In the absence of Csy4, in contrast, the *FAA1* site was edited to nearly 100%, while the *ADE2* editing was sharply reduced. In our multitron, the *ADE2* donor/gRNA was in the first position, suggesting that Csy4 processing is required on the 3’ end, adjacent to the gRNA scaffold, but dispensable on the 5’ end, adjacent to the msr.

It is preferable to minimize the donor/gRNA unit for practical reasons of construction, just as in the prokaryotic version. Therefore, in a parallel to the prokaryotic msd array multitron, we engineered a eukaryotic msd/gRNA array multitron, transferring the msr to a distinct transcript to reduce editing unit size and avoid long direct repeats (**Fig 5c**). This enabled construction of multitrons of arbitrary size using efficient one-step golden gate cloning. The msd encoding the donor remains fused to its matched gRNA, while a trans msr is able to function as a primer to create the RT-Donor internally (**Fig 5d**). We tested versions of this eukaryotic arrayed msd/gRNA multitron to precisely edit two, three, or five non-adjacent sites simultaneously (**Fig 5e-g**). In each case, all targeted sites were edited at a rate that increased over time. Thus, the arrayed msd multitron with trans msr is a generalizable strategy for multiplexing edits within a genome.

## DISCUSSION

This work demonstrates the construction, optimization, and use of multitrons for multiplexed precise editing within genomes of prokaryotic and eukaryotic cells. Final versions make use of donor-encoding retron msd arrays. Critically, we engineered the msd array format by optimizing not only for editing efficiency, but also for enabling practical cellular and molecular workflows. The compact multitron form is compatible with single-plasmid designs, one-step golden gate assembly, and plasmid removal in prokaryotic cells. These features should permit widespread adoption of the multitron editing approach.

We demonstrate simultaneous editing of up to five sites, with replacements of up to 8 base pairs per site, and deletions of up to 100 bases. This approach builds on previous work using oligonucleotides for MAGE by enabling efficient multisite editing without repeated transformations and by enabling a user to specify distinct combinations of donors per cell rather than relying on the random segregation of electroporated oligos. Multitrons enable a wider range of precise mutations than multiplexed base editors, and a more compact and simplified form than multiplexed prime editors.

We demonstrate proof-of-concept uses in molecular recording, genetic element minimization, and metabolic engineering. Future development will likely push the scale of multitrons both in the number of simultaneous mutations and the diversity of combinatorial mutations using libraries targeting two or more sites.

## METHODS

Biological replicates were taken from distinct samples, not the same sample measured repeatedly.

### Plasmid Construction

All the plasmids used in this work are listed in **Supplemental Table 2**. Furthermore, all the RT-donors and the oligonucleotides containing the desired mutations for the editing experiments are listed in **Supplemental Table 3.**

#### E. coli

To clone additional 70 bp donors in a single msd, pSLS.492^29^ plasmid containing a *rpoB* donor was used as backbone. To clone a donor upstream of the *rpoB* donor, a 60 bp reverse oligo annealing (25bp) with the 5’ region of the msd and containing 35 bp of the new donor, and a 60 bp forward oligo annealing (25bp) with the 5’ end of *rpoB* donor and harboring the other half of the new donor were used. To clone a donor downstream of the *rpoB* donor, a 60 bp forward oligo annealing (25b) with the 3’ region of the msd and containing 35 bp of the new donor, and a 60 bp reverse oligo annealing (25bp) with the 3’ end of *rpoB* donor and harboring the other half of the new donor were used. After a 30 cycles PCR reaction with Q5 hot-start high-fidelity polymerase (NEB) following recommended vendor protocol, a KLD reaction (NEB) was carried out to self-ligate the plasmid encoding an additional donor. To add a third donor in a single msd, the mentioned cycle was repeated again.

To construct the plasmids harboring the retron arrays in their different architectures the pCDF-DUET-1 vector (Novagen) was used as a backbone. A parental plasmid (pAGD159; **Supplemental Table** 2) containing a whole ncRNA with a *gyrA* donor downstream of the first T7 promoter, and Eco1-RT downstream of the second T7 promoter was constructed. To assess whole ncRNA retron arrays, the ncRNA harboring the rpoB donor from pSLS.492 was amplified and cloned upstream and downstream of the *gyrA*-containing ncRNA by Gibson Assembly. To construct the plasmids containing the msd array, firstly, the msr was deleted from pAGD159 and subsequently cloned between the second T7 promoter and Eco1-RT using a Gibson Assembly approach. Finally, the msd harboring the rpoB donor from pSLS.492 was amplified and cloned upstream and downstream of the *gyrA*-containing ncRNA by Gibson Assembly. To test if the msd array could act as a single transcript unit independent of the msr region, a T7 terminator was cloned between the msd array and the second T7 promoter.

To construct multitrons containing more than 2 arrayed msd a one-pot Golden Gate cloning approach was used. Firstly, a plasmid containing a sfGFP stuffer flanked by two inverted BsaI (type IIS restriction enzyme) target sites were cloned in the place of the msd Array generating pAGD236 **(see Figure 4c for reference).** Editing units, based on a msd with a donor were order as gBlocks (IDT) flanked by inverted BsaI target sites and compatible nucleotide overhangs to clone them in tandem. The Golden Gate protocol was carried out in 20uL reactions as follows: 1 uL pAGD236, 5uL of each gBlock (3uL for 5x msd arrays), 1.5uL BsaI (NEB), 2uL T4 DNA ligase Buffer, 0.5 uL T4 DNA ligase (NEB). Depending on the complexite of the reaction (more number of editing units) The reaction consists on 30 or 60 cycles (depending on the complexity) of 5 min at 16°C and 5 min at 37°C and a final cycle of 10 min at 60°C.

To optimize multitrons for metabolic engineering, the retron cassette (ncRNA and RT) from pSLS.492 was cloned into pORTMAGE-Ec1^33^ upstream of the CspRecT gene **(Supplemental Figure 2)**. RBS optimization of Eco1 RT and CspRecT genes were carried out using primers that contain the optimized RBS and self-ligating the plasmids using KLD reaction mix. Finally, recombineering operon was cloned into pKD-46^35^ backbone to obtain the parental temperature-sensitive multitron plasmid (pAGD248). Multitron msd array architectutre with the sfGFP stuffer flanked by two inverted BsaI described previously was cloned into pAGD248 generating pAGD335. The above-mentioned golden gate reaction was used to clone gBlocks containing the required donors into the pAGD335 backbone to generate the multitrons versions used in **Fig 4**.

#### S. cerevisiae

To assess whether Csy4 could enable the processing of editrons and retron msd/Cas9 gRNA units for genome editing, pSCL390, a derivative of pZS.157 (Addgene #114454), was generated with a yeast codon-optimized P2A-Csy4 CDS gblock (IDT) cloned downstream of the SpCas9 CDS by Gibson Assembly.

To compare the genome editing efficiencies of ribozyme-processed editrons to Csy4-processed editrons, pSCL.396, a derivative of pSCL.39 (Addgene #184973), was generated with the 5’ Hammerhead ribozyme and 3’ HDV ribozyme replaced by Csy4 recognition sites by amplification of the editron and backbone from pSCL.39 and assembled via Gibson Assembly.

To assess whether Csy4 could enable the processing of arrayed editrons, we generated pSCL.391, a derivative of pSCL.39 where a second editron, targeting the *S. cerevisiae* FAA1 locus was added on the 3’ end of the ADE2-targeting editron by Gibson Assembly. The cassette thus consists of two editrons, separated by a Csy4 recognition site, and flanked by a Hammerhead ribozyme and a HDV ribozyme on the 5’ and 3’ of the expression cassette, respectively.

To construct plasmids for the expression of retron msd arrays, first, a Golden Gate compatible entry vector, pSCL.452 was generated that carries the Gal7 promoter and terminator, alongside a cassette for expression of the retron msr from a Pol III SNR52 promoter. pSCL.452 is a derivative of a derivative of pSCL.39, generated by Gibson Assembly of the pSCL.39 backbone, amplified to replace the recombitron with inverted PaqCI sites for Golden Gate assembly, with a gblock (IDT) encoding pSNR52p-msr-SUP4t.

Next, plasmids carrying retron msd arrays for the editing of multiple loci in the yeast genome were generated by Golden Gate cloning of pre PaqCI-digested pSCL.452 with gBlocks (IDT) that encoded a PaqCI cutsite, a retron msd-encoded donor and paired gRNA for editing, a Csy4 recognition sequence, and a PaqCI cutsite **(Fig 5c)**. gBlocks were ordered with compatible nucleotide overhangs to enable random cloning of all combinations of gblocks into the entry plasmid, after PaqCI digestion. We ordered gblocks to edit the ADE2, FAA1, TRP2, SGS1 and CAN1 loci. These were cloned into the PaqCI-digested pSCL.452 backbone by Golden Gate cloning, yielding plasmids pSCL.473 (editors for ADE2, FAA1), pSCL.475 (editors for ADE2, CAN1 and FAA1) and pSCL.672 (editors for ADE2, FAA1, TRP2, SGS1 and CAN1).

### Strains and Growth Conditions

All bacterial and yeast strains are listed in Supplemental Table 4.

#### Bacterial Strains

The *E. coli* strains used in this study were DH5α (New England Biolabs) for cloning purposes, bMS.346 (DE3) for retron recombineering assays. Bacteria were grown in LB medium (10 g/l tryptone, 5 g/l yeast extract, 5 g/l NaCl). Antibiotics were added as required (carbenicillin, spectinomycin, kanamycin and chloramphenicol).

#### Yeast Strains

All yeast strains were created by LiAc/SS carrier DNA/PEG transformation^43^ of BY4742^26^. Strains for evaluating the effect of Csy4 on genome editing efficiency were created by BY4742 integration of plasmids pZS.157 (Addgene #114454) or pSCL.390. The plasmids were KpnI-linearized and inserted into the genome by homologous recombination into the *HIS3* locus. Transformants were isolated on SC-HIS plates.

### Bacterial Recombineering expression and analysis

In experiments using multitrons to edit bacterial genomes, the retron cassette encoded in a pET- 21 (+) plasmid (Novagen) and the CspRecT and mutLE32K in the plasmid pORTMAGE-Ec1^33^ were overexpressed using 1 mM IPTG, 1 mM m-toluic acid and 0.2% arabinose for 16 h with shaking at 37°C. For the molecular recording assay **(Fig 1g)**, a control without mtol and different concentration of the inducer, ranging from 0,005 mM to 0,1 mM, were added. To engineer the lycopene metabolic pathway **(Fig 4)**, bMS.346 electrocompetent cells containing pAC-LYC^42^ plasmid, were transformed with different multitron plasmid versions **(Supplemental Table 2 and 3).** and growth for 16 h at 30°C. Single colonies from the transformation plate were inoculated into 500uL of LB in triplicates in 1mL deep-well plates and incubated at 30°C for 24 h with vigorous shaking to prevent the cells from settling. A 1:1000 dilution of the cultures were passaged into LB 1% galactose and incubated at 30°C for 24 h with vigorous shaking. This last step was repeated for a total of 72h of editing.

After the different type of assays carried out in this study, a volume of 25 μl of culture was collected, mixed with 25 μl of water and incubated at 95°C for 10 min. A volume of 1 μl of this boiled culture was used as a template in 30-μl reactions with primers flanking the edit site, which additionally contained adapters for Illumina sequencing preparation (**Supplemental Table 5**). These amplicons were indexed and sequenced on an Illumina MiSeq instrument and processed with custom Python software to quantify the percentage of precisely edited genomes.

### Yeast editing expression and analysis

The parental strains (–Csy4: HIS3::pZS.157; +Csy4: HIS3::pSCL390) were transformed with variants of the editron expression cassettes by LiAc/SS carrier DNA/PEG transformation. Single colonies from the transformation plate were inoculated into 500uL of SC-HIS-URA 2% raffinose in triplicates in 1mL deep-well plates and incubated at 30°C for 24 h with vigorous shaking to prevent the cells from settling. Cultures were passaged into SC-HIS-URA 2% galactose and incubated at 30°C for 24 h with vigorous shaking. This was repeated once more for experiments meant to compare the genome editing efficiencies of ribozyme-processed editrons to Csy4-processed editrons, for a total of 48h of editing; and four more times for experiments meant to assess whether arrays of retron msds could be used to edit multiple loci in the yeast genome, for a total of 120h of editing. At each timepoint of galactose-induced editing, a 250uL aliquot of the cultures was harvested, pelleted and washed with water, and prepped for deep sequencing of the loci of interest.

Samples were prepped for deep sequencing of the edited loci as described previously^29^. Briefly, genomic DNA was extracted by (1) resuspending the cell pellets in 120uL of lysis buffer (100 mM EDTA pH 8, 50 mM Tris-HCl pH 8, 2% SDS) and heating them to 95 °C for 15 min; (2) cooling the lysate on ice and adding 60uL of protein precipitation buffer (7.5 M ammonium acetate), then inverting gently and placing samples at -20°C for 10min; (3) centrifugation of the samples at maximum speed for 2mins (or until a clear supernatant forms) and collecting the supernatant (∼100uL) in new 1.5mL tubes; (4) precipitating the nucleic acids by adding equal parts of ice-cold isopropanol to the samples, mixing the samples thoroughly and incubating the mix at -20°C for 10min (or overnight for higher yield), followed by pelleting by centrifugation at maximum speed for 2min; (5) washing the pellet twice with 200 µl of ice-cold 70% ethanol, followed by air-drying it; and (6) resuspending the pellet in 40 µl of water. 0.5uL of gDNA was used as template in 20-µl PCR reactions with primers flanking the edit site in of the target locus, which additionally contained adapters for Illumina sequencing preparation (**Supplemental Table 5**). Importantly, the primers do not bind to the retron msd donor sequence. These amplicons were indexed and sequenced on an Illumina MiSeq instrument and processed with custom Python software to quantify the percentage of precise edits using the retron derived RT-DNA template.

### Colorimetric screen and assay for lycopene production

After cycle 3 (72h) of the metabolic engineering assay, cells from the edited bMS.346 populations using diferrent multitrons were plated on LB-chloramphenicol agar plates and grown for 1 day at 30°C and 2 days more in darkness and at room temperature to produce red colonies. Per edited population with a multitron, plates containing around 10^3^ colonies were screened by visual inspection searching for increased red colour intensity. A total of 84 colonies (12 isolates from each multitron version and 12 from the control) were selected for lycopene quantification. These isolated colonies were grown into 1 mL LB-chloramphenicol in 1 mL deep-well plates for 24 h at 37°C to cure multitron plasmid. For lycopene extraction, 1 ml of cells were centrifuged at 16,000g for 30 s, the supernatant was removed and the cell pellet was resuspended with 1 mL water. Cells were re-centrifuged at 16,000g for 30 s, the supernatant was removed and the cells were resuspended in 200 ml acetone and incubated in the dark for 15 min at 55 °C with intermittent vortexing. The mixture was centrifuged at 16,000g for 1 min and the supernatant containing the lycopene was transferred to 96 white/clear bottom plate. Absorbance at 470 nm of the extracted lycopene solution was measured using a spectrophotometer to determine the lycopene content. Lycopene yield of the different colonies from each was calculated by normalizing the times of lycopene production against the control. Cells coming from different clusted of lycopene production were re-striked in LB-cloramphenicol agar plates grown for 24h at 30°C and for another 48h at room temperature. Between 3 and 8 colonies from each re-striking were selected to quantify the lycopene production following the described protocol and for Sanger sequencing across the *dxs/idi* targets.

### Data Availability

All data supporting the findings of this study are available within the article and its supplementary information, or will be made available from the authors upon request. Sequencing data associated with this study will be available on NCBI SRA prior to peer-reviewed publication.

### Code Availability

Custom code to process or analyze data from this study will be made available on GitHub prior to peer-reviewed publication.

## Acknowledgements

Work was supported by funding from the National Science Foundation (MCB 2137692), the National Institute of Biomedical Imaging and Bioengineering (R21EB031393), the National Institute of General Medical Sciences (1DP2GM140917), and the UCSF Program for Breakthrough Biomedical Research. S.L.S. is a Chan Zuckerberg Biohub – San Francisco Investigator and acknowledges additional funding support from the L.K. Whittier Foundation and the Pew Biomedical Scholars Program. A.G.-D. was supported by the California Institute of Regenerative Medicine (CIRM) scholar program. S.C.L. was supported by a Berkeley Fellowship for Graduate Study.

## Author Contributions

S.L.S., A.G.-D. and S.C.L. conceived the study. A.G.-D., S.C.L., M.R.M. and C.B.F. performed the experiments; A.G.-D., S.C.L. and S.L.S. designed the work, analyzed the data and wrote the manuscript.

## CORRESPONDING AUTHOR

Correspondence to Seth L. Shipman.

---

**Supplementary Figure 1.**
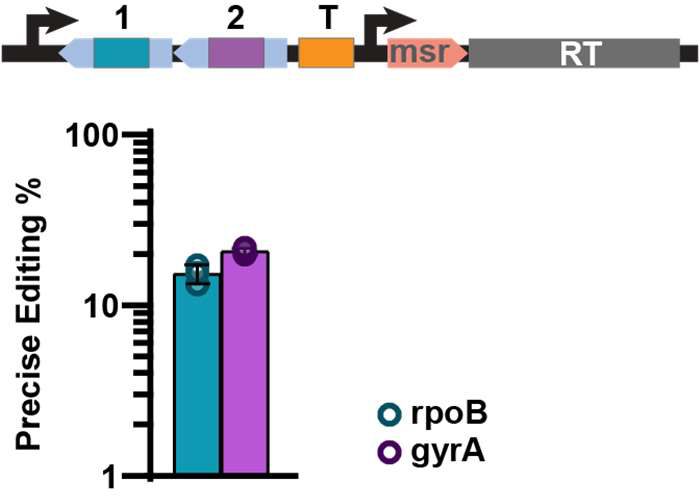
Trans msr multitron architecture enables precise genome editing. Top: Schematic of retron recombineering using an msd array with a single msr sequence in trans including a terminator (T) between the msd array and msr. Bottom: quantification of precise editing rates for precise editing of *rpoB* or *gyrA* simultaneously by Illumina sequencing after 24h of editing. Circles show each of the three biological replicates, bars are mean ±SD.

**Supplementary Figure 2.**
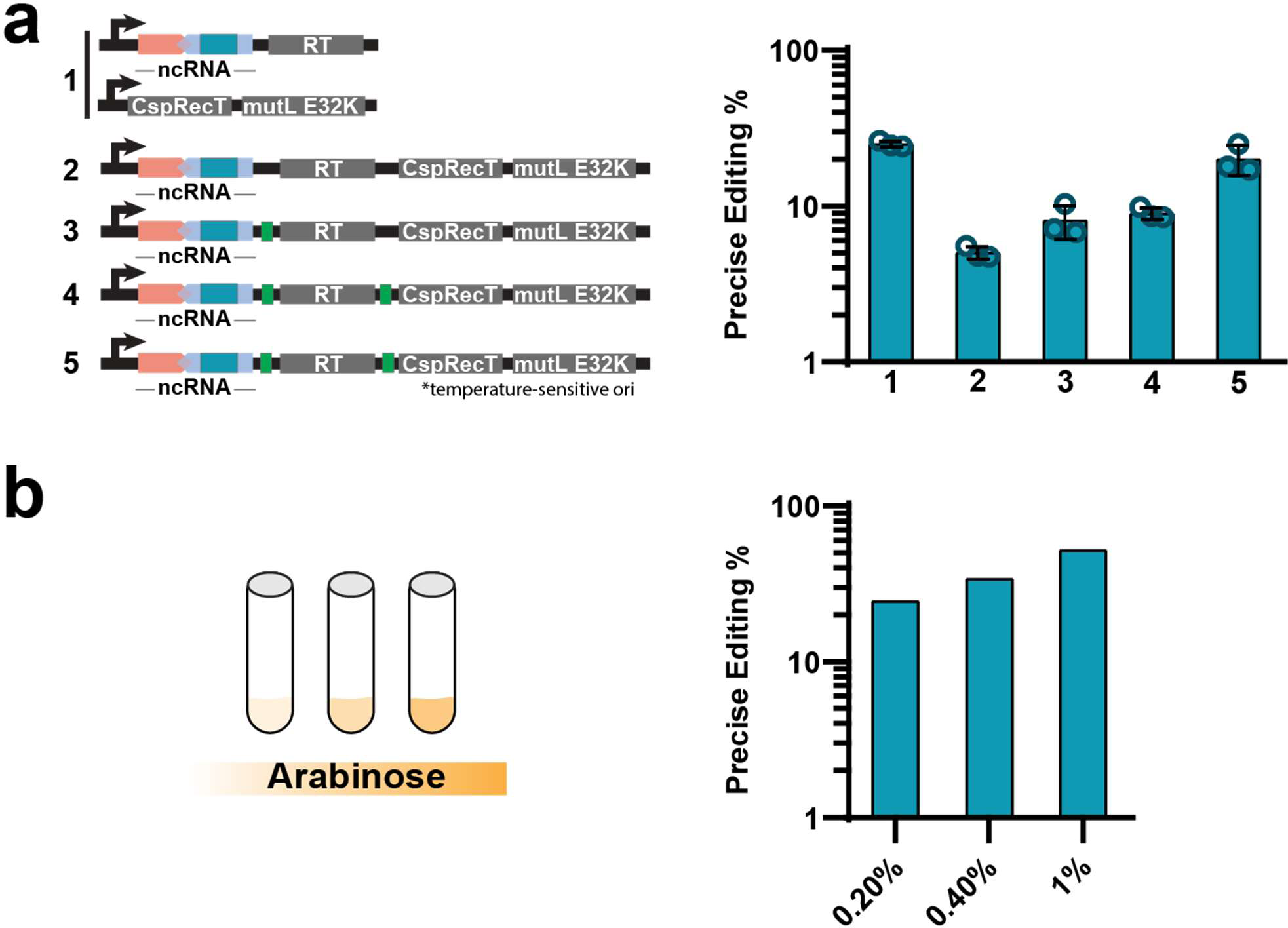
Optimization of retron recombineering plasmids for metabolic engineering. **a.** Left: schematic of the different retron operon architectures tested. ncRNA with donor (orange and blue), genes required (grey) and optimized ribosome binding sites (RBS) regions (green) are indicated Right: quantification of rates for precise *rpoB* editing, circles show each of the three biological replicates, bars are mean ±SD. **b.** Left: schematic of arabinose titration to optimize retron recombineering. Right: quantification of precise editing rates for *rpoB* using different concentrations of arabinose (n=1). All data was quantified using Illumina MiSeq after 24h of editing.

